# Biological sex determines skeletal muscle atrophy in response to cortical TDP-43 pathology

**DOI:** 10.1101/2024.09.01.610721

**Authors:** G Lorenzo Odierna, Laura A Reale, Tracey C Dickson, Catherine A Blizzard

## Abstract

**Background:** Amyotrophic lateral sclerosis (ALS) is a fatal and incurable neurodegenerative condition. In ALS, wasting of skeletal muscle causes weakness, paralysis and ultimately, death due to respiratory failure. Diagnosis of ALS is a long process and delays in diagnosis are common, which impedes rapid provision of patient care and treatment. Additional tools or methodologies that improve early detection might help overcome the diagnostic delays and enhance survival and quality of life for people with ALS. In this study, we used a transgenic mouse model to create a detailed catalogue of skeletal muscle wasting with the goal of finding muscles that can be examined to enhance early diagnosis of ALS.

**Methods:** Cortical pathology was induced by crossing CaMKIIa-tTA and tetO-hTDP-43^ΔNLS^ transgenic mice (ΔNLS). Transgenic expression was induced at 30-days postnatal via removal of doxycycline diet. Mice were aged to 15-, 20-, 30- and 45-days post transgene induction. Microdissection was applied to isolate 22 individual hindlimb muscles for measurement of weight. Both males and females were used at all timepoints.

**Results:** We found that male and female ΔNLS mice exhibited hindlimb skeletal muscle atrophy relative to controls. Multiply innervated muscles, also known as series-fibered muscles, were especially vulnerable to atrophy. The strongest predictor of the atrophic response across all hindlimb muscles was the extent to which any individual muscle was larger in males than females, known also as sexual dimorphism. In males, muscles that are usually larger in males compared to females experienced the most atrophy. Conversely, in females, muscles that are usually of similar size between males and females experienced the most atrophy. Segregating muscles based on whether they were more affected in males or females revealed that hip extensors, knee flexors, knee extensors, ankle dorsiflexors and ankle evertors were more affected in males. Hip adductors, hip rotators, hip flexors and ankle plantarflexors were more affected in females.

**Conclusions:** Our results demonstrate that the difference in the size of skeletal muscles in males compared to females is the most powerful predictor of muscle atrophy in response to dying forward pathology. This indicates that sex is a strong determinant of skeletal muscle vulnerability in ALS. Our results provide new insights into determinants of skeletal muscle atrophy and may help inform selection of muscles for diagnostic testing of ALS patients.

## Introduction

Amyotrophic lateral sclerosis (ALS) is a fatal, progressive neurodegenerative condition that aggressively destroys the motor system. It is characterised by dysfunction of both upper and lower motor neurons, presenting as weakness and atrophic wasting of voluntary skeletal muscles. The main cause of death in people with ALS is respiratory failure due to paralysis of the muscles that control breathing [1], which usually occurs within 2-5 years of disease onset [2]. Incidence of ALS, presentation of symptoms, and disease progression have all been linked to biological sex [3, 4]. The global sex ratio for ALS incidence is skewed towards men [5, 6] but certain subtypes of the disease occur more frequently in women or men. Bulbar onset ALS, for example, more commonly affects women [7, 8] whereas limb onset ALS more commonly affects men [9, 10]. Diagnosis of ALS is also sensitive to sex. Although most studies report that diagnosis takes 10-16 months from when symptoms first appear [11–13], men with spinal onset ALS are at higher risk of delayed diagnosis compared to women and are more than twice as likely to receive a misdiagnosis [14, 15]. Given the short survival for people with ALS, there is a strong incentive to understand the factors that contribute to the heterogeneity of the disease and the challenges associated with delivering rapid, accurate diagnosis.

Skeletal muscle currently provides one of the most important avenues for diagnosing ALS, primarily because muscle dysfunction directly underlies symptom presentation. Muscle biopsies and motor unit studies indicate that the earliest symptoms of skeletal muscle dysfunction in ALS are driven by neuromuscular denervation and reinnervation [16–22]. Symptoms caused by this denervation-reinnervation cycle, such as fasciculations, can be detected via electrodiagnostic neurological examination [23]. Electromyography and needle exam typically play an instrumental role in providing an ALS diagnosis but are time consuming and spatially limited to assessing single muscles. This means that selection of which muscles to assess is an important step in the diagnostic process. Indeed, most criteria used to reach a diagnostic conclusion of ALS require signs of motor neuron dysfunction in multiple body regions, detectable by either muscle weakness or electromyographic abnormalities [24, 25]. In the absence of observable muscle atrophy or weakness, knowing which muscles to target for electromyographic assessment would accelerate diagnosis of ALS. A recent study by Fukushima et al. (2022) applied machine learning to muscle ultrasonography data collected from early-stage ALS patients and revealed that not all muscles hold equal diagnostic value [26]. This is in line with previous observations that some muscles, such as the tongue, provide special diagnostic utility [27–29]. With increasing recognition that there exist muscles with high diagnostic value, additional work mapping the skeletomuscular system in ALS holds the potential to minimise the diagnostic delay for patients.

One of the factors that differentiates ALS from other neuromotor disorders is the involvement of the motor cortex and upper motor neurons. Clinical studies have revealed that people with ALS display signatures of increased excitability of the motor cortex [30–32]. Importantly, this hyperexcitability is an early biosignature of ALS; it can be detected in people with family history of disease, before onset of symptoms and before clinically detectable lower motor neuron dysfunction [30, 33]. These data provide support for the ‘dying forward’ hypothesis, which claims ALS pathogenesis initiates in the brain and spreads anterogradely throughout the corticomotor system [34]. One of the most striking influences of dying forward pathology is that muscles receiving stronger cortical drive are selectively vulnerable in ALS, leading to the ‘split phenomena’. The most well-known of these is the split hand sign, which is caused by selective wasting of the thenar muscles over the hypothenar and is relatively specific to ALS over other neuromotor disorders [35].

We have recently used a transgenic mouse model to study how dying forward pathology spreads throughout the motor system. It permits temporally inducible expression of transgenes with a specific enrichment in the cortex and reduced expression in the spinal cord [36], including absence from lower motor neurons [37]. Driving expression of an NLS-mutated human TDP-43 protein (hTDP-43^ΔNLS^) this way causes cytoplasmic TDP-43 accumulation [36], which recapitulates the cytoplasmic cellular TDP-43 inclusions found in upwards of 97% of people with ALS [38–40]. We have found that pyramidal neurons in layer 5 of the motor cortex display hyperexcitability in these mice as early as 20-days post expression [37, 41, 42]. Following this, at 30-days post expression, the ventral horn of the spinal cord undergoes synaptic remodelling [37]. Because hTDP-43^ΔNLS^ expression is largely absent from the ventral horn our findings suggest that the changes induced in the cortex, where hTDP-43^ΔNLS^ expression is strongest, cascade down the corticomotor system in a ‘dying forward’-like manner. In this study, we capitalised on the well-established timeline and cortical pathology of the hTDP-43^ΔNLS^ model with the goal of identifying skeletal muscle targets with high diagnostic value to ALS.

## Methods

### Mouse husbandry

All mouse experiments were conducted with the approval of The University of Tasmania Animal Ethics Committee (approval number A24715) in compliance with the 2013 Australian Code of Practice for the Care and Use of Animals for Scientific Purposes. In this study, CaMKIIa-tTA and tetO-hTDP-43^ΔNLS^ were used to generate bigenic mice with an active tetracycline transactivator system. Monogenic mice (CaMKIIa-tTA or tetO-hTDP-43^ΔNLS^) were used as controls. Stock animals were originally sourced from The Jackson Laboratory (CaMKIIa-tTA RRID JAX:003010 and tetO-hTDP-43^ΔNLS^ RRID JAX:014650) and then backcrossed for 10 generations to a C57BL/6J background. Breeding mice and their offspring were maintained on a stock of chow containing 200 mg/kg doxycycline. For experimental mice, dox chow was switched to standard chow between P28-P33 until they were euthanised. Experimental mice were group housed (2-5 per cage) based on sex in individually ventilated cages, with *ad libitum* access to food and water. Upon reaching their designated experimental end points, mice were humanely euthanised by intraperitoneal injection of pentobarbital using a terminal dose of 200 mg/kg. Both sexes of mice were used for all experiments in this study.

### Muscle isolation via microdissection

Following humane killing via pentobarbital overdose, mice were transcardially perfused with 4% paraformaldehyde in 0.1 M phosphate buffered saline (PBS) and fixed overnight. We aimed to isolate and weigh as many individual hindlimb muscles as we could confidently dissect, including at least one muscle per major functional group. Functional grouping was based on definitions by Charles et al., 2016. We reiterate here the recognition of the authors of that study that many muscles are bi-articular and that the functional grouping is based on maximum moment arm values. We were able to successfully isolate muscles involved in hip extension (*semimembranosus*, SM; *semitendinosus,* ST; *biceps femoris anterior*, BFA; *biceps femoris posterior*, BFP; *caudofemoralis*, CA), hip adduction (*adductor longus*, AL; *adductor magnus*, AM; *gracilis anterior*, GA; *gracilis posterior*, GP), hip flexion (*pectineus*; PECT), hip rotation (*quadratus femoris*, QF), knee extension (*rectus femoris*, RF), knee flexion (*popliteus*, POP), ankle dorsiflexion (*tibialis anterior*, TA: *extensor digitorum longus*, EDL), ankle plantarflexion (*gastrocnemius*, GAS; *plantaris*, PL; *soleus*, SOL; *flexor digitorum longus*, FDL; *tibialis posterior*, TP) and ankle eversion (*peroneus longus*, PL; *peroneus tertius*, PT). Care was taken to ensure that tendons were severed at similar locations across all dissected muscles and direct muscular attachment points to bone were separated as closely to the bone as possible. Dissected muscles were stored at 4°C in 0.1M phosphate buffered saline containing 0.1% sodium azide.

### Western blots

Following humane killing via pentobarbital overdose, mice were transcardially perfused with ice cold 0.1 M PBS. Brain and muscle tissue samples were rapidly extracted, snap frozen in liquid nitrogen and stored at -80°C. Protein was extracted from tissue samples using modified Radio Immunoprecipitation Assay (RIPA) buffer containing: 150 mM NaCl, 50 mM Tris pH 8, 5 mM EDTA, 1% NP-40 and 0.5% Sodium Deoxycholate, with protease inhibitor tablet (Roche, catalogue #04693132001). Tissue samples were manually cut into small pieces on dry ice using a cold scalpel and then incubated in RIPA buffer (20 µl per mg of tissue) overnight at 4°C on a shaker. Samples were centrifuged at 13,000 rpm for 20 minutes at 4°C. The supernatant was removed, and protein concentration was determined using a Bradford protein assay.

Lysates were separated using polyacrylamide gel electrophoresis. Proteins were denatured by heating samples at 80°C for 10 minutes in NuPAGE LDS sample buffer (Thermo Fisher Scientific, catalogue #NP0007) containing 5% β-mercaptoethanol. 20µg of protein per sample was loaded onto NuPAGE 4- 12% Bis-Tris gels (Thermo Fisher Scientific, catalogue #NP0336BOX). Electrophoresis was performed in NuPAGE MES SDS running buffer (Thermo Fisher Scientific, catalogue #NP002) and transfer to PVDF membrane was performed in NuPAGE transfer buffer (Thermo Fisher Scientific, catalogue # NP0006). Membranes were blocked for 1 hour in Tris Buffered Saline (TBS) containing 0.01% Tween- 20 and 5% non-fat milk. Membranes were incubated in primary antibodies in TBS containing 0.01% Tween-20 and 5% non-fat milk overnight at 4°C. Membranes were washed in TBS, then incubated in secondary antibodies in TBS containing 0.01% Tween-20 and 5% non-fat milk for 1 hour. Immobilon chemiluminescent HRP substrate was used for detection via an Amersham I600 imager. Membranes were exposed for 2minutes to maximise any possible weak signals in muscle tissue lysates. Membranes were stripped and re-probed for a loading control.

The following antibodies were used in this study: mouse anti-TDP-43 (human-specific, 1:10000, ProteinTech, Cat# 60019-2, RRID# AB_2200520), rabbit anti-GAPDH (1:20000, Millipore, Cat# ABS16, RRID# AB_10806772), goat anti-mouse IgG HRP conjugate (1:40000, Dako, Cat# P0447, RRID# AB_2617137).

### Statistical tests and analyses

In this study, we define atrophy based on the average weight of muscles in CaMKIIa-hTDP-43^ΔNLS^ mice relative to monogenic controls. In all instances where the value “% Atrophy” is referred to, the value was calculated according to: 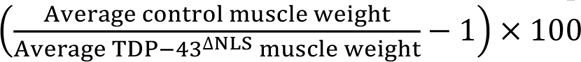

In this study, we define sexual dimorphism based on the average weight of male muscles relative to the average weight of female muscles. In all instances where the value “% Sexual dimorphism” is referred to, the value was calculated according to 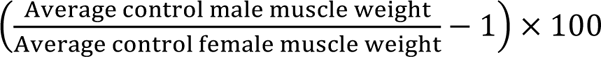

To test for statistically significant differences between groups, a two-way analysis of variance (ANOVA) was used with Tukey’s multiple comparisons test. Correlations were assessed by performing a simple linear regression and statistical significance of calculated slopes from baseline (slope=0) was determined using a sum-of-squares F test. Significance threshold was set at *p*<0.05. All analyses were performed in GraphPad Prism (version 10.1.2). Relevant details pertaining to tests used, error bars and N numbers can be found in figure legends.

## Results

### CaMKIIa-dependent conditional hTDP-43^ΔNLS^ expression causes body weight loss not attributable to slower growth or muscular hTDP-43^ΔNLS^ expression

Post-developmental expression of hTDP-43^ΔNLS^ in the cortex induces loss of body weight and functional motor behavioural defects [36, 44]. In this study, we tracked and analysed body weight of male and female mice separately. A significant reduction in body weight was detectable in CaMKIIa-hTDP- 43^ΔNLS^ (hereon “ΔNLS”) relative to control male mice by 20-days post expression and ΔNLS and relative to control female mice by 30-days post expression (Figure 1A). This decrease in body weight persisted until 60-days post expression in both males and females. Thus, although reduced body weight occurs in both sexes of ΔNLS mice compared to controls, it is detectable at least 10 days earlier in males than in females.

**Figure 1.**
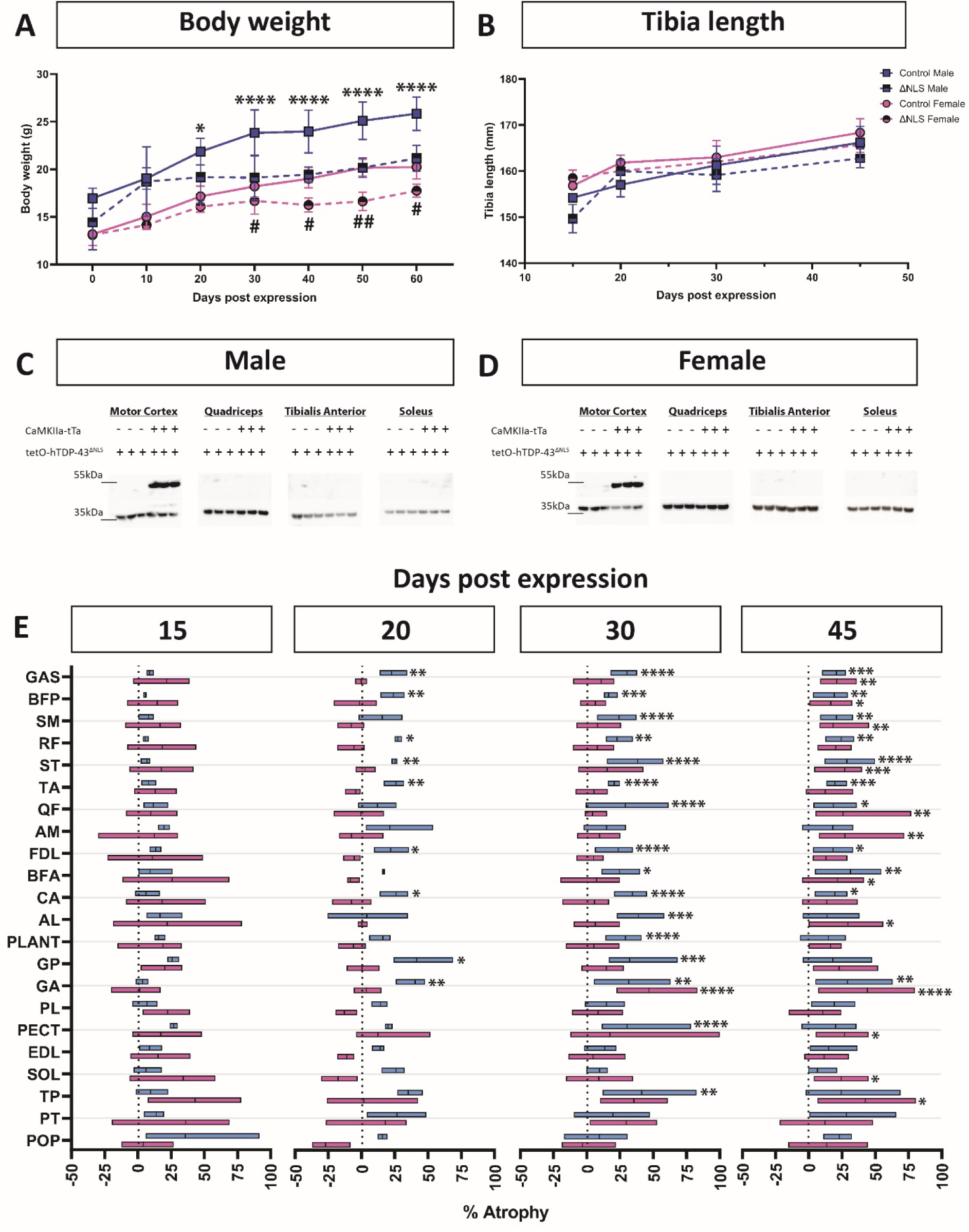
Hindlimb skeletal muscles experience atrophy in CaMKIIa-hTDP-43^ΔNLS^ mice relative to controls. (**A**) Body weight of male and female CaMKIIa-hTDP-43^ΔNLS^ (ΔNLS) mice compared to controls (monogenic CaMKIIa-tTA or tetO-hTDP-43^ΔNLS^). Transgenic expression was induced at 30- days postnatal (P30), and measurements were made every 10-days post expression. Groups analysed using a two-way ANOVA with Tukey’s multiple comparisons test, asterisks (*) indicate significance for male ΔNLS mice relative to male controls and hashes (^#^) indicate significance for female ΔNLS mice relative to female controls. \**p*<0.05, \*\*\*\**p*<0.0001, ^##^*p*<0.01. N numbers for control male at each timepoint post expression are: 0, N=6; 10, N=6, 20, N=6; 30, N=11; 40, N=5; 50, N=5, 60, N=5. N numbers for control female at each timepoint post expression are: 0, N=6; 10, N=10; 20, N=10; 30, N=13; 40, N=3; 50, N=3; 60, N=3. N numbers for ΔNLS male at each timepoint post expression are: 0, N=3; 10, N=6; 20, N=6; 30, N=14; 40, N=8; 50, N=8; 60, N=8. N numbers for ΔNLS female at each timepoint post expression are: 0, N=7; 10, N=7; 20, N=7; 30, N=14; 40, N=8; 50, N=8; 60, N=8. (**B**) Tibia length of male and female ΔNLS mice compared to controls following induction of transgene expression at P30. Groups analysed using a two-way ANOVA with Tukey’s multiple comparisons test. N numbers for control male at each timepoint post expression are: 15, N=5; 20, N=3; 30, N=4; 45, N=6. N numbers for control female at each timepoint post expression are: 15, N=4; 20, N=4; 30, N=3; 45, N=5. N numbers for ΔNLS male at each timepoint post expression are: 15, N=3; 20, N=4; 30, N=8; 45, N=9. N numbers for ΔNLS female at each timepoint post expression are: 15, N=5; 20, N=3; 30, N=4; 45, N=7. (**C**, **D**) Western blots of male (**C**) and female (**D**) tissue lysates probed using antibodies against human TDP-43 (top, ∼43 kDa) and GAPDH (bottom, ∼37 kDa). Lysates were generated from motor cortex, quadriceps femoris muscles (Quadriceps), tibialis anterior (TA) muscle and soleus (SOL) muscle. Genotype indicated via presence (+) or absence (-) of CaMKIIa-tTA or tetO-hTDP-43^ΔNLS^ transgenes. (**E**) Muscle weight data, available in Table S1, expressed as % Atrophy, calculated as follows: [(average control muscle weight/average ΔNLS muscle weight)-1]*100. Data visualised as a box plot where horizontal box length reflects range and vertical line indicates mean. Blue boxes are male, pink boxes are female. Muscles ordered based on size from largest (top) to smallest (bottom). The difference between control and ΔNLS was tested using a two-way ANOVA with Tukey’s multiple comparisons test for each muscle separately (sex and days post expression as independent variables). Asterisks denote a significant difference between control and ΔNLS for males and females separately. \**p*<0.05, \*\**p*<0.01, \*\*\**p*<0.001, \*\*\*\**p*<0.0001. The difference between % Atrophy between males and females was not tested. GAS, gastrocnemius; BFP, biceps femoris posterior; SM, semimembranosus; RF, rectus femoris; ST, semitendinosus; TA, tibialis anterior; QF, quadratus femoris; AM, adductor magnus; FDL, flexor digitorum longus; BFA, biceps femoris anterior; CA, caudofemoralis; AL, adductor longus; PLANT, plantaris; GP, gracilis posterior; GA, gracilis anterior; PL, peroneus longus; PECT, pectineus; EDL, extensor digitorum longus; SOL, soleus; TP, tibialis posterior; PT, peroneus tertius; POP, popliteus. N numbers for all muscles (barring exceptions) of control male at each timepoint post expression are: 15, N=4; 20, N=4; 30, N=9; 45, N=4. Exceptions were AM (20, N=3), QF (20, N=3; 45, N=3), PECT (20, N=3; 45, N=3), POP (45, N=3), PL (45, N=3), PT (15, N=3; 45, N=3), TP (45, N=3), FDL (45, N=3). N numbers for all muscles (barring exceptions) of control female at each timepoint post expression are: 15, N=6; 20, N=5; 30, N=4; 45, N=6. Exceptions were GP (20, N=4), GA (20, N=4), QF (20, N=4), POP (20, N=4). N numbers for all muscles of ΔNLS male at each timepoint post expression are: 15, N=3; 20, N=3; 30, N=5; 45, N=7. N numbers for all muscles (barring exceptions) of ΔNLS female at each timepoint post expression are: 15, N=7; 20, N=6; 30, N=8; 45, N=9. Exceptions were AM (20, N=5), AL (20, N=5), QF (30, N=7), PECT (20, N=5), PT (15, N=6), TP (15, N=6; 45, N=8).

We reasoned that decreased body weight may be driven by a slower rate of growth in ΔNLS mice or muscle atrophy. To test the first possibility, we measured tibia length seeing as this directly correlates with whole animal growth [45]. Tibia length was assessed at 4 separate time points: 15-, 20-, 30- and 45-days post expression. There was no statistically significant difference between ΔNLS and control mice in both males and females across all time points (Figure 1B). The decrease in body weight in these mice therefore cannot be explained by slower animal growth, suggesting that it may instead be caused by skeletal muscle atrophy.

In considering the possibility of skeletal muscle atrophy in ΔNLS mice we wanted to rule out the possibility of local hTDP-43^ΔNLS^ expression in muscle. The presence of TDP-43 pathology in muscle is associated with degeneration [46] and therefore muscle-specific expression might provide a simple explanation for the observed loss of body weight. We performed western blots using whole muscle lysates generated from hindlimb muscles collected at 30-days post expression including the quadriceps femoris, tibialis anterior and soleus. As a positive control, we assessed expression of hTDP-43 in the motor cortex, where expression is known to be high. Consistent with previous reports, we found strong expression of hTDP-43 in the motor cortex of both male and female mice (Figure 1 C, D). Conversely, all hindlimb skeletal muscles assessed did not display any evidence of any hTDP-43 expression in both sexes (Figure 1 C, D). The lack of evidence for hTDP-43 expression in the quadriceps femoris, tibialis anterior and soleus confirms the specificity of this model as a forebrain-specific driver of pathology, meaning that any effects in skeletal muscle must occur due to spread of pathology.

### Hindlimb muscles atrophy earlier in males than females and in a heterogenous pattern

Our previous work has demonstrated that lower motor neurons do not express hTDP-43 in ΔNLS mice but that the spinal cord undergoes synaptic remodelling consistent with a ‘dying forward’-like spread of pathogenic changes from the cortex [37]. Having confirmed that hTDP-43 is also not expressed in skeletal muscle in this mouse model, we interrogated how anterograde spread of pathology impacted muscle outcomes in the periphery. To do this, we isolated and weighed individual hindlimb muscles of ΔNLS and control mice at 15-, 20-, 30- and 45-days post expression. We were able to successfully isolate 22 muscles (muscles listed in methods), representing ∼60% of all reported skeletal muscles in the C57BL/6 mouse hindlimb and covering 100% of its major functional groups [43, 47].

After weighing all muscles (data in Table S1), we calculated the average atrophy for each one in ΔNLS relative to controls (see methods for details). We found that muscles in ΔNLS mice were atrophied relative to control and that sex strongly influenced this across all the timepoints we investigated. We also found that there was substantial heterogeneity in the extent of atrophy in ΔNLS mice relative to controls at the level of each individual muscle (Figure 1E). At 15-days post expression, muscles in ΔNLS mice were not atrophied relative to controls in both males and females. At 20-days post expression, no muscles were atrophied in females whereas ∼40% of the assessed muscles displayed atrophy in males. At 30-days post expression, only one muscle was atrophied in females (the GA), compared to ∼70% of the assessed muscles in males. Lastly, at 45-days post expression, ∼55% of the assessed muscles were atrophied in females compared 50% in muscles. Thus, in this model, the induction of hindlimb skeletal muscle atrophy occurs later in females than males. Moreover, there is substantial heterogeneity in which muscles experience atrophy and the extent to which atrophy occurs amongst those that do.

### Atrophy does not correlate with the fiber type composition of skeletal muscles

One of the most striking characteristics of the atrophic response of skeletal muscle in ΔNLS mice was the heterogeneity between individual muscles and between biological sexes (Figure 1E). The sampling breadth of our hindlimb muscle weight data allowed us to interrogate the source of this variance. To begin, we tested whether muscle atrophied based on their fiber typing. Fast fatigable motor units involving glycolytic muscle fibers (type II) are selectively vulnerable in ALS [48–51]. In order to test whether fiber type explained the differences in atrophy we observed, we focused on six muscles which have had their fiber type composition previously mapped: the EDL, TA, GAS, PLANT, SOL and QF [52–54]. The inclusion of the QF in our study is unique since it provides another predominantly oxidative muscle to compare to the SOL. The QF is, in fact, comprised of more oxidative fibers than the SOL [54], meaning it should experience as much or even more protection from pathology. When we compared the muscle weight data for the QF and the SOL, however, we found that this was not the case. Although the SOL in male mice was not affected at all timepoints, the QF was atrophied at both 30- and 45-days post expression (Figure 2A, B). In female mice, both the SOL and the QF were atrophied at 45-days post expression (Figure 2A, B). To further visualise how these muscles compared to known glycolytic muscles, we generated scatterplots for atrophy against fiber type composition (% type I fibers). This reinforced our original observations and performing a linear regression analysis on the data revealed poor R^2^ values with slopes that were not statistically different from baseline (Table 1). Our data thus show that fiber type composition is not a good predictor of atrophy in ΔNLS mice.

**Figure 2.**
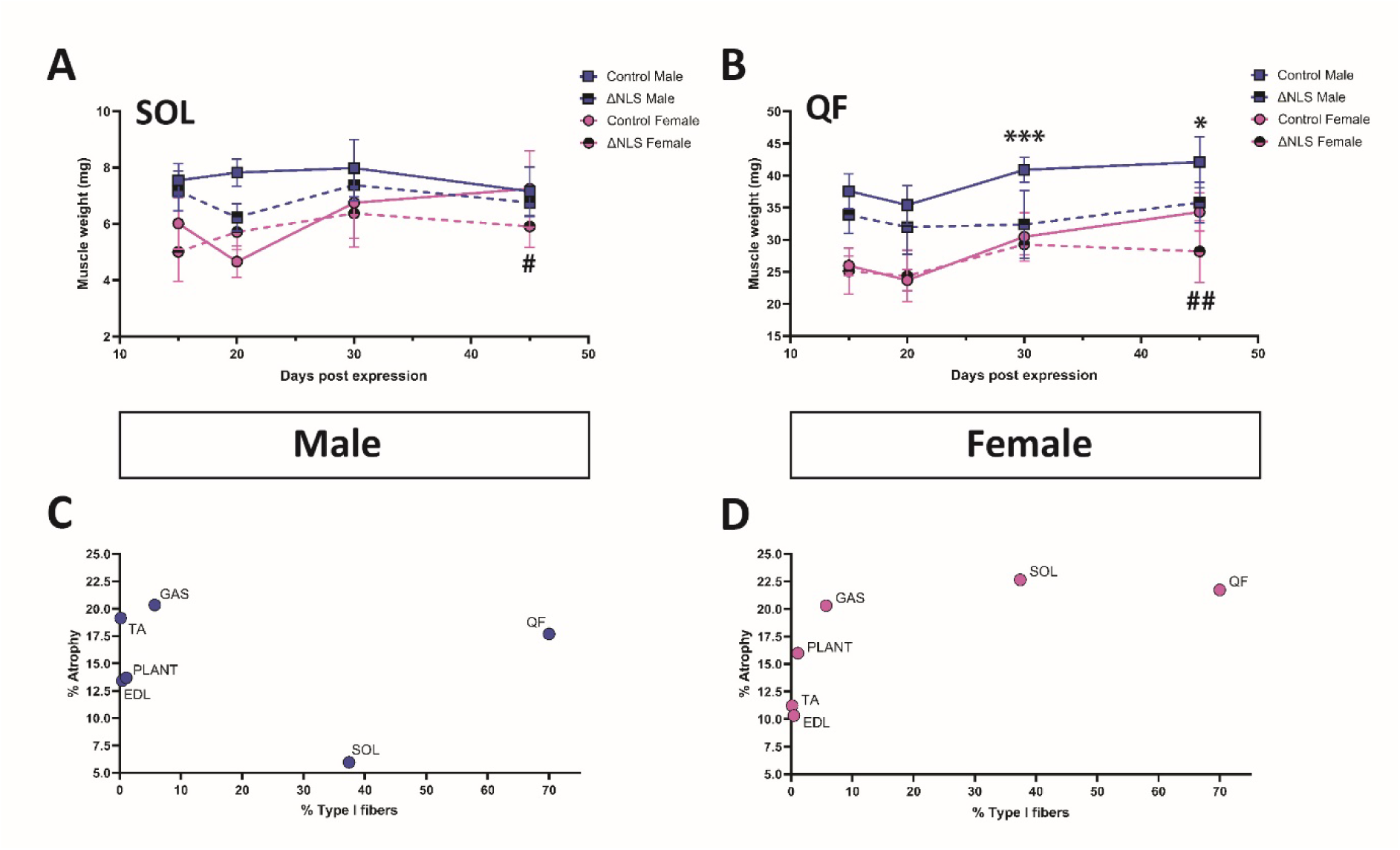
Fiber type composition does not predict atrophy in CaMKIIa-hTDP-43^ΔNLS^ mice. (**A**, **B**) Weight measurement of soleus (SOL; **A**) and quadratus femoris (QF; **B**) in control male, control female, CaMKIIa-hTDP-43^ΔNLS^ (ΔNLS) male and ΔNLS female mice. Groups analysed using a two- way ANOVA with Tukey’s multiple comparisons test, asterisks (*) indicate significance for male ΔNLS mice relative to male controls and hashes (#) indicate significance for female ΔNLS mice relative to female controls. \**p*<0.05, \*\*\*\**p*<0.0001, ^#^*p*<0.05, ^##^*p*<0.01. N numbers for control male at each timepoint post expression are: 15, N=4; 20, N=4 (QF, N=3); 30, N=9; 45, N=5 (QF, N=3). N numbers for control female at each timepoint post expression are: 15, N=6; 20, N=5 (QF, N=4); 30, N=4; 45, N=6. N numbers for ΔNLS male at each timepoint post expression are: 15, N=3; 20, N=3; 30, N=6 (SOL, N=5); 45, N=7. N numbers for ΔNLS female at each timepoint post expression are: 15, N=7; 20, N=6; 30, N=8 (QF, N=7); 45, N=9. Error bars show standard deviation. (**C**, **D**) Scatterplot of % Atrophy against reported composition of oxidative fibers (% Type I fibers) in males (**C**) and females (**D**) at 45-days post expression. % Atrophy was calculated as follows: [(average control muscle weight/average ΔNLS muscle weight)-1]*100. Fiber type composition data sourced from Augusto et al. (2004), DeNies et al. (2014) and Hitomi et al. (2005). Each point represents the average % Atrophy per muscle (indicated in graph) across all mice. SOL, soleus; PLANT, plantaris; GAS, gastrocnemius; TA, tibialis anterior; EDL, extensor digitorum longus; QF, quadratus femoris.

**Table 1.** Linear regression analyses of cortical hTDP-43^ΔNLS^ expression-induced atrophy against properties of all hindlimb muscles.

|  | Days post expression | Male |  |  | Female |  |  |
| --- | --- | --- | --- | --- | --- | --- | --- |
|  |  | Slope | R<sup>2</sup> | p-value | Slope | R<sup>2</sup> | p-value |
| Pennation angle<sup>†</sup> | 15 | -0.8±0.4 | 0.2 | 0.1 | -0.004±1.3 | 9×10<sup>-7</sup> | 0.9 |
|  | 20 | 0.2±1.2 | 0.003 | 0.9 | 0.7±1.6 | 0.02 | 0.7 |
|  | 30 | 3.3±1 | 0.5 | 0.01* | -0.7±1.8 | 0.02 | 0.7 |
|  | 45 | 1.1±0.9 | 0.1 | 0.3 | 0.7±1.6 | 0.02 | 0.7 |
| Oxidative fiber type composition<sup>‡</sup> | 15 | -0.009±0.05 | 0.006 | 0.9 | -0.02±0.1 | 0.01 | 0.8 |
|  | 20 | -0.07±0.1 | 0.1 | 0.5 | -0.01±0.1 | 0.002 | 0.9 |
|  | 30 | -0.008±0.1 | 0.0007 | 0.9 | -0.007±0.04 | 0.007 | 0.9 |
|  | 45 | -0.03±0.09 | 0.03 | 0.8 | 0.1±0.06 | 0.5 | 0.1 |
| Weight | 15 | -0.09±0.04 | 0.2 | 0.09 | 0.02±0.07 | 0.006 | 0.7 |
|  | 20 | 0.02±0.06 | 0.003 | 0.8 | 0.06±0.09 | 0.02 | 0.5 |
|  | 30 | 0.03±0.06 | 0.01 | 0.6 | -0.04±0.07 | 0.02 | 0.6 |
|  | 45 | 0.03±0.03 | 0.05 | 0.3 | -0.001±0.06 | 0.0005 | 0.9 |
| Sexual size dimorphism | 15 | 0.5±0.09 | 0.6 | <0.0001* | -0.2±0.1 | 0.2 | 0.06 |
|  | 20 | 0.4±0.2 | 0.2 | 0.03* | -0.4±0.2 | 0.3 | 0.02* |
|  | 30 | 0.8±0.2 | 0.5 | 0.0009* | -0.6±0.2 | 0.4 | 0.002* |
|  | 45 | 0.4±0.08 | 0.5 | 0.0002* | -0.4±0.1 | 0.3 | 0.02* |

\* Statistical deviation of slope from baseline (0) tested using extra sum-of-squares F test with significance threshold set to 0.05.
† Values for pennation angle sourced from Charles et al. (2016). Only pennate muscles were used for this analysis: Pectineus, rectus femoris, tibialis anterior, extensor digitorum longus, gastrocnemius, plantaris, soleus, flexor digitorum longus, tibialis posterior, peroneus longus and peroneus tertius.
‡ Values for oxidative fiber type composition sourced from Augusto et al. (2004), DeNies et al. (2014) and Hitomi et al. (2005). Muscles used for this analysis include the quadratus femoris, gastrocnemius, plantaris, soleus, tibialis anterior and extensor digitorum longus.
Note, semitendinosus and gracilis anterior muscles were not included for any linear regression analyses at all since these muscles have an series-fibered arrangement and are uniquely affected in this mouse model (Figure 4C, D).

### Musculoskeletal anatomy and architecture do not explain heterogeneity of the atrophic responses

After finding that fiber type did not explain the pattern of atrophy in ΔNLS mice, we turned to other possible explanations such as anatomical location or muscle fiber architecture. We segregated the muscles based on whether they acted on the hip (Figure 3A), knee (Figure 3B) or ankle (Figure 3C). We found that splitting the muscles this way did not reveal any meaningful patterns; muscles that act on the hip, knee and ankle were affected equally in males (Figure 3D) and females (Figure 3F) across all timepoints. Next, we split muscles into two groups based on whether their muscle fibers had a pennate or non-pennate organisation. Splitting muscles this way revealed that both pennate and non- pennate muscles were equally affected at all timepoints in males (Figure 3E) and females (Figure 3G). To further test the effect of pennation we performed a linear correlation analysis between atrophy and pennation angle (for pennate muscles only; data sourced from Charles et al., 2016). The analyses revealed generally poor R^2^ values and that the slope of the regression was not discriminable from baseline for the majority of timepoints in males and females (Table 1). The analysis for males at 30- days post expression was an exception to this since it revealed that pennation angle explained 50% of the variance in atrophy with a slope that was statistically different from baseline (Table 1). However, the largely poor performance of pennation angle as predictor of atrophy in males and females at all other timepoints did not provide us with confidence that it serves as a determinant of selective vulnerability. Thus, anatomical location and fiber pennation architecture do not explain the atrophic response of skeletal hindlimb muscles in ΔNLS mice.

**Figure 3.**
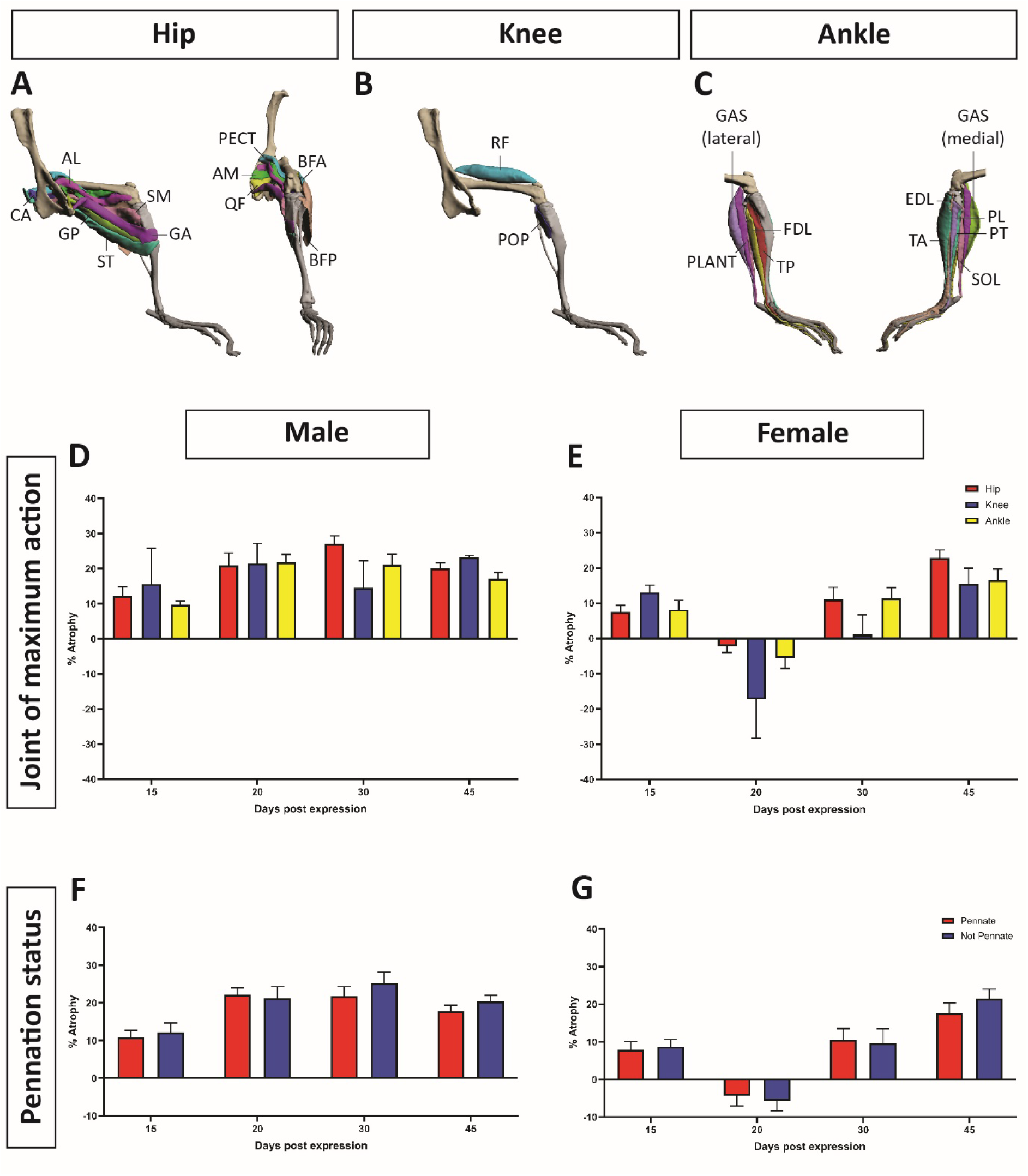
Muscles that act on the hip, knee and ankle are equally affected, as are pennate and not pennate muscles. (**A**-**C**) Visualisation of muscles used in this study, separated by the joint on which they act maximally. Functional categorisation based on maximal moment arm values in Charles et al. 2016. 3D renders of hindlimb muscles adapted from Charles et al., 2016 (Creative Commons Attribution License, CC-BY 4.0). The hip muscles we isolated are shown in panel **A** and include the biceps femoris posterior (BFP), biceps femoris anterior (BFA), semitendinosus (ST), semimembranosus (SM), caudofemoralis (CA), adductor magnus (AM), adductor longus (AL), gracilis anterior (GA), gracilis posterior (GP), quadratus femoris (QF) and pectineus (PECT). The knee muscles we isolated are shown in panel **B** and include the rectus femoris (RF) and popliteus (POP). The ankle muscles we isolated are shown in panel **C** and include the gastrocnemius (GAS), plantaris (PLANT), soleus (SOL), flexor digitorum longus (FDL), tibialis posterior (TP), tibialis anterior (TA), extensor digitorum longus (EDL), peroneus longus (PL) and peroneus tertius (PT). (**D**, **E**) Muscle atrophy of male (**D**) and female (**E**) CaMKIIa-hTDP-43^ΔNLS^ (ΔNLS) mice compared based on joint of maximum action. % Atrophy was calculated as follows: [(average control muscle weight/average ΔNLS muscle weight)-1]*100. Groups analysed using a two-way ANOVA with Tukey’s multiple comparisons test. N numbers reflected number of muscles segregated into each category. Hip N=11, knee N=2, ankle N=9. Error bars show standard error of the mean. (**F**, **G**) Muscle atrophy of male (**F**) and female (**G**) mice compared based on pennation status. Groups analysed using a two-way ANOVA with Tukey’s multiple comparisons test. N numbers reflected number of muscles segregated into each category. Pennate N=11, not Pennate N= 11. Error bars show standard error of the mean.

### Series-fibered muscles are selectively vulnerably to atrophy in both males and females

One less considered property of muscles in ALS research is their serialisation status. Series-fibered muscles differ from parallel-fibered muscles in that instead of being comprised of individual muscle fibers which run contiguously from tendon to tendon (in parallel), they are formed by multiple independent sets of muscle fibers which terminate intramuscularly and consequently are innervated by multiple endplate bands (in series) [55]. The mouse hindlimb contains two series-fibered muscles: the GA and the ST. The GA is particularly valuable when studying series-fibered muscles because it is one of two muscles, alongside the GP, that make up the gracilis in mice. Both the GA and the GP are similarly sized, have similar attachment points and play similar functional roles. This means that the biggest difference between them is their serialisation status. When we compared the muscle weight data for the GA and the GP, we found that there was a noticeable difference in their atrophic responses. Notably, the GA was much more strongly affected than the GP both in magnitude and across time (Figure 4A, B). This was especially obvious in female mice, for which the GA was the only muscle to have experienced atrophy by 30-days post expression (Figure 4B). To test whether this observation held true more generally, we segregated all hindlimb muscles based on their serialisation status. We found that series-fibered muscles were more atrophied than parallel-fibered muscles in both male and female mice in a statistically significant manner. Series-fibered muscles consistently experienced ∼33% more atrophy than parallel-fibered muscles at 20-, 30- and 45-days post expression in males (Figure 4C). In females, series-fibered muscles experienced ∼66% more atrophy than parallel-fibered muscles at 30- days post expression and ∼42% more at 45-days post expression (Figure 4D). Thus, muscles with a series-fibered arrangement are selectively vulnerable to atrophy in response to cortical pathology.

**Figure 4.**
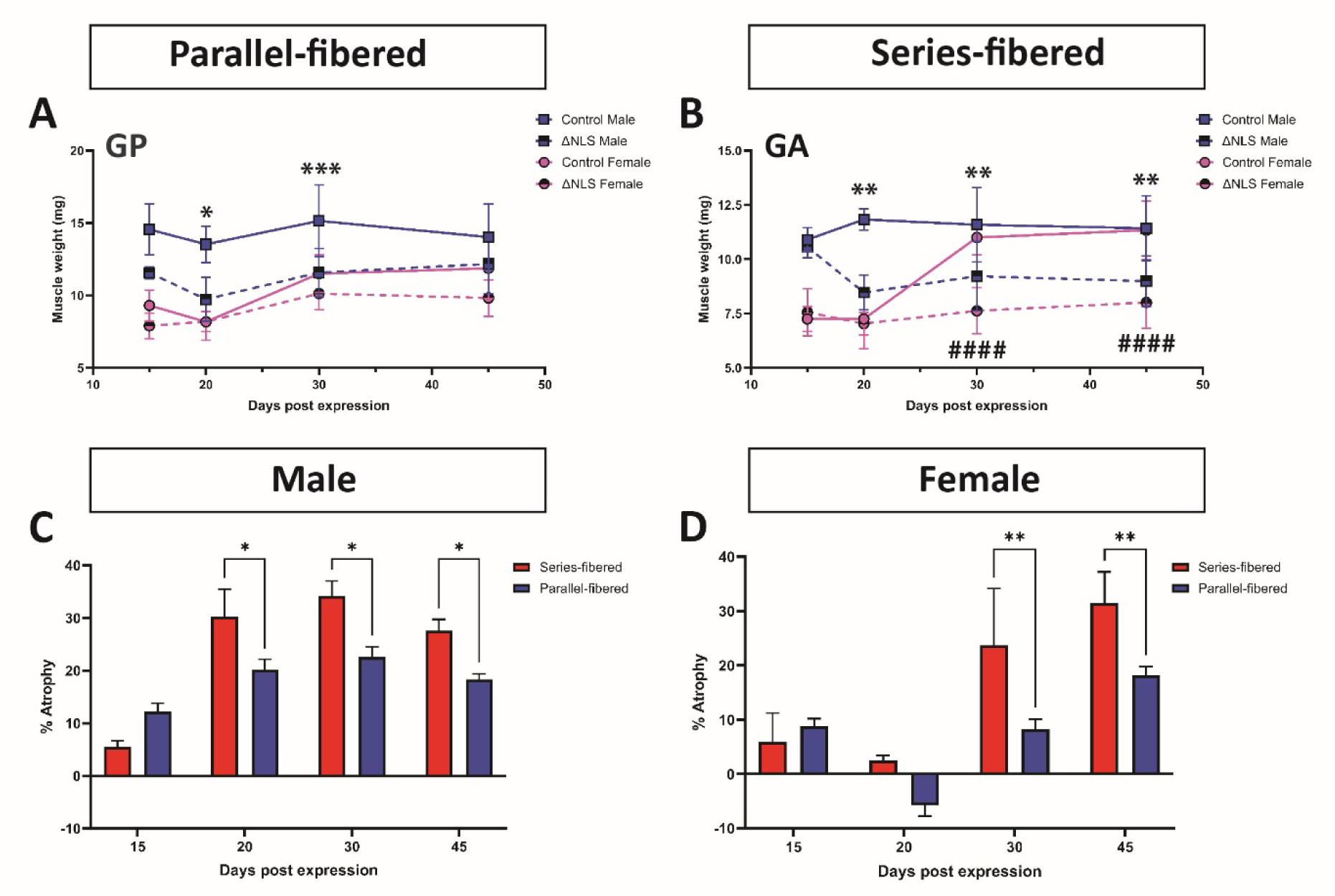
Series-fibered muscles are selectively vulnerable relative to parallel-fibered muscles. (**A**, **B**) Weight measurement of gracilis posterior (GP; **A**) and gracilis anterior (GA; **B**) in control male, control female, CaMKIIa-hTDP-43^ΔNLS^ (ΔNLS) male and ΔNLS female mice. Groups analysed using a two-way ANOVA with Tukey’s multiple comparisons test, asterisks (*) indicate significance for male ΔNLS mice relative to male controls and hashes (#) indicate significance for female ΔNLS mice relative to female controls. \**p*<0.05, \*\**p*<0.01, \*\*\**p*<0.001, ^####^*p*<0.0001. N numbers for control male at each timepoint post expression are: 15, N=4; 20, N=4; 30, N=9; 45, N=5 (GP, N=4). N numbers for control female at each timepoint post expression are: 15, N=6; 20, N=4; 30, N=4; 45, N=6. N numbers for ΔNLS male at each timepoint post expression are: 15, N=3; 20, N=3; 30, N=5; 45, N=7. N numbers for ΔNLS female at each timepoint post expression are: 15, N=7; 20, N=6; 30, N=8; 45, N=9. Error bars show standard deviation. (**D**, **E**) Muscle atrophy of male (**D**) and female (**E**) ΔNLS mice compared based on serialisation status. % Atrophy was calculated as follows: [(average control muscle weight/average ΔNLS muscle weight)-1]*100. Groups analysed using a two-way ANOVA with Tukey’s multiple comparisons test. \**p*<0.05, \*\**p*<0.01. N numbers reflected number of muscles segregated into each category. Series-fibered N=3 (ST separated into ST-P and ST-D for this analysis), parallel-fibered N=20. Error bars show standard error of the mean.

### Sexual dimorphism is the strongest predictor of skeletal muscle atrophy in CaMKIIa-hTDP- 43^ΔNLS^ mice

Despite having found that series-fibered arrangement was a significant determinant of atrophy, we could not explain what determined the heterogeneity in atrophy for the remaining muscles. We plotted weight (based on values from control mice in this study) against atrophy and calculated the extent of correlation by performing a linear regression but the results from this analysis indicated that muscle size was not correlated with atrophy in ΔNLS mice (R^2^ values ≤0.2 at all the timepoints in both sexes, no *p*-values crossed threshold; Table 1). Next, we calculated the difference in size between male and female muscles by dividing the average weight of each muscle in male controls by the average weight of each muscle in female controls and expressing this as a percentage (see methods for details; all values outlined in Table S2). We found that this ratio, which is formally known as ‘sexual dimorphism’ [56], varied significantly across hindlimb muscles almost forming a continuum (Figure 5A). When we performed a linear regression analysis between sexual dimorphism and atrophy, we found that it was a powerful predictor of muscle atrophy in both males and females at multiple timepoints. In males, sexual dimorphism was positively associated with atrophy with a slope that had a statistically significant deviation from baseline at 15- though to 45-days post expression. The R^2^ values ranged from 0.2-0.6 across all timepoints (Table 1) and by 45-days post expression, sexual dimorphism explained 50% of the variance in muscle atrophy (Figure 5B). In females, this relationship was inverted such that sexual dimorphism was negatively associated with atrophy. The calculated slope of the linear regression had a statistically significant deviation from baseline from 20- to 45-days post expression. The R^2^ values ranged from 0.2-0.4 across all timepoints (Table 1) and by 45-days post expression, sexual dimorphism explained 30% of the variance in muscle atrophy (Figure 5C). Taken together, these data demonstrate that sexual dimorphism is the greatest predictor of muscle atrophy in ΔNLS mice. Further, the relationship between atrophy and sexual dimorphism is strongly influenced by biological sex. In males, the most strongly affected muscles are those that are bigger in males compared to females (Figure 5B). Conversely, in females, the most strongly affected muscles are those that are of equal size in males and females (Figure 5C).

**Figure 5.**
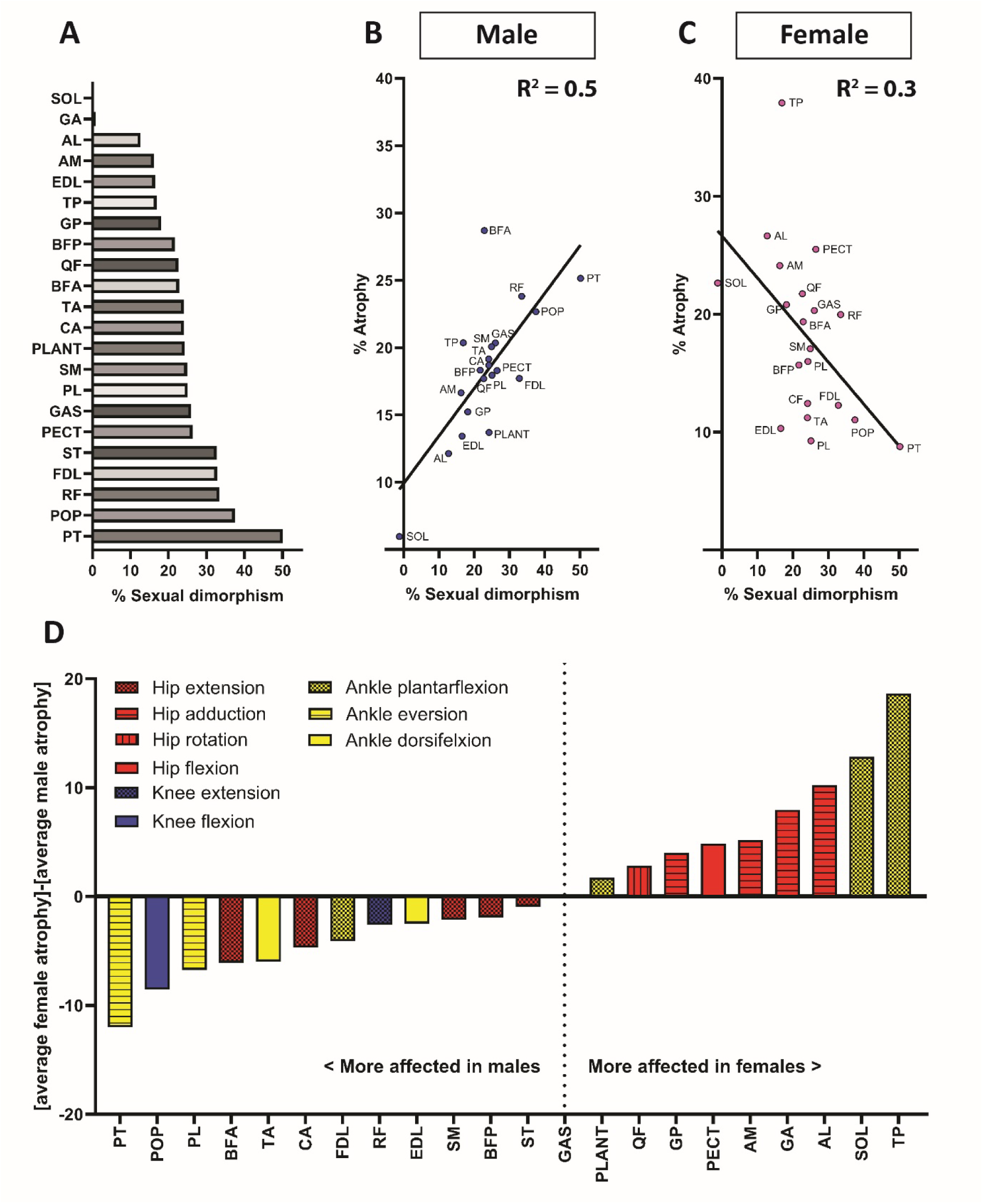
Sexual dimorphism predicts atrophy in CaMKIIa-hTDP-43^ΔNLS^ mice. (**A**) Bar plot of sexual dimorphism per muscle, calculated as follows: [(Average control male muscle weight/Average control female muscle weight)-1]*100. Muscles are displayed in order of % Sexual dimorphism, from lowest (top) to highest (bottom). SOL, soleus; GA, gracilis anterior; AL, adductor longus; AM, adductor magnus; EDL, extensor digitorum longus; TP, tibialis posterior; GP, gracilis posterior; BFP, biceps femoris posterior; QF, quadratus femoris; BFA, biceps femoris anterior; TA, tibialis anterior; CA, caudofemoralis; PLANT, plantaris; SM, semimembranosus; PL, peroneus longus; GAS, gastrocnemius; PECT, pectineus; ST, semitendinosus; FDL, flexor digitorum longus; RF, rectus femoris; POP, popliteus; PT, peroneus tertius. (**B**, **C**) Scatterplots of % Atrophy against % Sexual dimorphism in males (**B**) and females (**C**) at 45-days post expression. % Atrophy was calculated as follows: [(average control muscle weight/average ΔNLS muscle weight)-1]*100. Each point represents the average % Atrophy per muscle (each muscle name indicated in graph) across all mice. Line of best fit from linear regression analysis overlayed on each scatterplot for male and female data (linear regression analysis results can be found in Table 1). Line slopes were significantly different to baseline (0) and each R^2^ value is overlayed. (**D**) Bar plot of the difference between average % Atrophy in males and females per muscle. Difference between average % Atrophy in males and females calculated using: [average female % Atrophy]-[average male % Atrophy]. Muscles are arranged in ascending order from those most affected in males to those most affected in females. Functional group of the muscle is indicated in the legend.

### Affected muscles segregate into different functional groups in males and females

Having found that sexual dimorphism was a strong predictor of atrophy, we next sought to test the extent to which biological sex contributed to atrophy in ΔNLS mice. We were particularly interested in the inversion of the relationship between sexual dimorphism and atrophy between males and females. One prediction from this result was that some muscles were more affected in males compared to females and *vice versa*. To investigate this, we revisited the weight data for all muscles across all timepoints. We noticed that there were clearly some muscles that were strongly impacted based on sex. The PL and PT, for example, were more affected in males than females at 45-days post expression whereas the TP and SOL were more strongly affected in females than males at this timepoint (Figure 1E). In order to test whether muscles segregated based on sex in any meaningful way, we ordered each muscle based on the difference between the atrophic response in males and females. When plotted this way, muscles segregated based on the major functional groups of the hindlimb (Figure 5D). For muscles that were more strongly affected in males than females, there was a noticeable bias towards muscles involved in hip extension, knee extension, knee flexion, ankle dorsiflexion and ankle eversion. Conversely, for muscles that were more strongly affected in females than males, there was a bias towards muscles involved in hip adduction, hip rotation, hip flexion and ankle plantar flexion. This data reinforces the idea that biological sex plays a significant role in determining the skeletal muscle response to cortical hTDP-43^ΔNLS^ pathology and suggests that this role is strongly connected to muscle function.

## Discussion

In this study, we explored the skeletal muscle response in CaMKIIa-hTDP-43^ΔNLS^ (ΔNLS) mice over time in both males and females separately. The ΔNLS mouse model is defined by cortically biased expression of human TDP-43 that harbours mutations in the NLS domain, causing cytoplasmic buildup of the protein [36]. One of the strengths of the model is its inducibility, which allows for the expression of the hTDP-43^ΔNLS^ to begin post-developmentally. Our previous work has shown that layer 5 pyramidal neurons of the motor cortex progressively develop hyperexcitability starting at 20-days post transgene induction [37, 41], followed by loss of excitatory synapses onto these cells [42, 57]. This is followed by synaptic remodelling in the ventral horn of the lumbar spinal cord, despite a lack of expression of the hTDP-43^ΔNLS^ in lower motor neurons [37]. The mice lose weight and develop both cognitive and motor behavioural deficits, which can be restored if transgene expression is repressed [44]. Here, we show that the loss of body weight in both male and female mice is attributable to skeletal muscle. The hTDP-43^ΔNLS^ protein is not expressed in skeletal muscle, demonstrating that the atrophy is not caused by TDP-43 pathology in muscle. Rather, it likely occurs as a consequence of a ‘dying forward’-like spread of changes originating in the cortex, where the hTDP-43^ΔNLS^ protein expression is highest. Our study therefore shows that cortical pathology can produce skeletal muscle atrophy. Intriguingly, male mice experienced muscle atrophy relative to controls as early as 20-days post expression, which coincides with the earliest detectable electrophysiological changes in the motor cortex of these mice [37]. The finding that the muscular atrophic response occurs concomitant to the cortical electrophysiological response indicates that a motor system-wide failure follows cortical expression of hTDP-43^ΔNLS^.

As part of our study, we performed at detailed analysis on muscle weight data, including ∼60% of all identified muscles in the C57BL/6 mouse hindlimb, covering 100% of all hindlimb functional groups. Our data spanned four separate time points post transgenic induction for both males and females. This dataset allowed us to identify sources of variation in the atrophic responses across all the muscles we collected. We found that multiply innervated muscles comprised of a series-fibered arrangement were more strongly affected in both male and female mice. This type of muscle fiber arrangement may thus represent a determinant of selective muscle vulnerability in ALS. In line with our findings, existing data suggests series-fibered muscles may be uniquely vulnerable in ALS patients. For example, the genioglossus has been identified as holding special diagnostic value in determining disease duration in ALS [58] and displays higher rates of fasciculations compared to other cranial muscles [59]. Of the four extrinsic muscles of the tongue, the genioglossus is the only one that shows evidence of having a series- fibered arrangement, due to the presence of multiple end plate bands [60]. Other series-fibered muscles such as the brachioradialis [61], gracilis and sartorius [62] have received less attention but may prove diagnostically useful for ALS. This is especially true when considering that they appear to be less affected in ALS mimics such as Kennedy’s disease and Hirayama’s disease [63] wherein the sartorius/gracilis and brachioradialis are spared, respectively [64, 65].

To further characterise determinants of atrophy, we performed correlations against several key muscle factors. Many factors were not associated with atrophy, including size, anatomical location and fiber type. The failure of fiber type to explain atrophy was unexpected, given the broadly reported selective vulnerability of type II fibers in ALS rodent models [48–51]. It is worth mentioning here that the ankle muscles which are most frequently studied in preclinical rodent research tended to follow previously reported trends for fiber type such that in male mice, the soleus (SOL) was much less atrophied than the extensor digitorum longus (EDL), tibialis anterior (TA) and gastrocnemius (GAS). Inclusion of the highly oxidative quadratus femoris (QF), which is a rarely studied hip rotator, revealed that oxidative fiber type alone does not provide a good explanation for the variance in atrophic responses we observed. Instead, the most powerful predictor of muscle atrophy was the extent to which any muscle was bigger in males compared to females, a ratio that is termed ‘sexual dimorphism’. By calculating this ratio for each muscle, we report here that sexual dimorphism across the hindlimb is highly variable. The extent of sexual dimorphism for each muscle was highly predictive of how strongly it atrophied in ΔNLS mice. Interestingly, the relationship between sexual dimorphism and atrophy was opposite in males and females. In males, muscles that were larger in males compared to females more experienced the most atrophy. Conversely, in females, muscles that were similarly sized in males and females experienced the most atrophy. To our knowledge, this is the first time that sexual dimorphism of skeletal muscles has been identified as a predictor of muscle atrophy in ALS. This result suggests that patient sex should be strongly considered during the early stages of diagnosing ALS and that men and women may possibly need to be assessed using separate criteria. It also contributes to a growing body of work that provides recognition of biological sex as a mechanistically relevant factor in ALS pathogenesis.

When we segregated muscles based on the difference in atrophic responses between males and females there were clear sex-based associations based on functional group. For hip muscles, hip extensors were more affected in males whereas hip adductors, flexor and rotators were more affected in females. For ankle muscles, ankle dorsiflexors and evertors were more affected in males whereas ankle plantarflexors were more affected in females. This dissociation based on functional grouping was somewhat reminiscent of the “split phenomena” in ALS. The split phenomena in ALS consist of the ‘split hand’[66], ‘split elbow’ [67] and ‘split leg’ [68] signs, which are characterised by dissociated/preferential wasting of muscles in the hands, arms and legs, respectively. Current hypotheses suggest that the dissociations observed in the split phenomena arise due to differences in cortical drive to some muscles over others [69, 70]. One theme that arises from research into the split phenomena is that discrepancies tend to arise with respect to which groups are specifically affected over others. For the split-elbow sign, Khalaf et al. (2019) reported relative sparing of the triceps brachii compared to the biceps brachii [67] whereas Liu et al. (2021) found the opposite - that the elbow flexors are less affected than the elbow extensors [71]. For the split-leg sign, Simon et al. (2015) reported relative sparing of the ankle dorsiflexors compared to the ankle plantarflexors [68] but this was not consistent with the indication of more strongly afflicted ankle dorsiflexors in other studies, more consistent with foot drop syndrome in ALS patients [71–74]. One possible explanation for these discrepancies is that the nature of the preferential wasting might vary depending on the cohort of patients recruited for each study. Although wiring of the corticomotor system in the murine and human nervous systems has many differences, our data suggests that some of the discrepancies in these patient studies may also possibly arise due to sex. The results from our work suggest that segregating by sex when studying split phenomena may prove insightful. Our data also suggest that the split phenomena may be present in muscles that act on the hip and that assessment of these muscles may provide a new avenue for diagnosing or assessing the progression of ALS.

## Conclusions

In this study, we set out to identify muscles with high diagnostic value to ALS. We used a dying forward- oriented model of ALS to correlate the extent of atrophy for individual muscles with a range of skeletal muscles properties. Our findings demonstrate that multiply innervated series-fibered muscles are selectively vulnerable to dying forward pathology. Further, we identify that the extent to which muscles differ in size between males and females is a powerful predictor of skeletal muscle atrophy. These results provide valuable insights into disease aetiology as well as new targets that can be used to enhance diagnosis of ALS.

## Acknowledgements

We would like to acknowledge the support of the MND Research Institute Australia, the National Health and Medical Research Council and the Tasmanian Medical Research Foundation. We thank the University of Tasmania Animal Services team and Veterinarians for their important contributions to this work.

## Data and resource availability

All relevant data and resources can be found within the article and its supplementary information.

## Author contributions

GLO: Conceptualization, Formal Analysis, Investigation, Data curation, Writing – original draft preparation, Writing – review and editing, Visualization. LAR: Investigation, Writing – review and editing. TCD: Resources, Writing – review and editing, Funding acquisition. CAB: Resources, Writing – review and editing, Funding acquisition.

---

**Table S1.** Muscle weight data for male and female control and CaMKIIa-hTDP-43^ΔNLS^ mice at 15-, 20-, 30- and 45-days post expression.

| **Muscle** | **Days post expression** | **Control male** | **ANLS male** | **Control female** | **ANLS female** |
| --- | --- | --- | --- | --- | --- |
| **GAS** | 15 | 112.2±7.8 | 103.2±2.7 | 79.3±4.0 | 66.2±9.5 |
|  | 20 | 111.8±10.5 | 91.5±7.6 | 75.3±1.7 | 75.5±2.3 |
|  | 30 | 128.9±8.9 | 99.0±5.6 | 91.1±6.0 | 83.1±10.2 |
|  | 45 | 124.8±7.3 | 103.7±5.3 | 99.0±8.7 | 82.3±6.2 |
| **BFP** | 15 | 99.9±5.5 | 94.7±1.2 | 71.6±4.3 | 63.4±9.1 |
|  | 20 | 103.8±10.6 | 84.1±6.2 | 70.4±6.2 | 72.4±9.8 |
|  | 30 | 119.6±7.3 | 102.8±3.3 | 88.2±9.1 | 83.3±6.3 |
|  | 45 | 119.2±3.0 | 100.7±9.4 | 97.9±9.7 | 84.7±7.5 |
| **SM** | 15 | 74.6±1.9 | 69.3±4.5 | 56.2±2.7 | 49.1±7.9 |
|  | 20 | 75.6±10.3 | 66.4±10.0 | 50.2±4.4 | 54.7±4.7 |
|  | 30 | 88.9±6.8 | 72.1±6.1 | 63.4±3.6 | 59.1±6.0 |
|  | 45 | 87.1±3.9 | 72.6±5.9 | 69.7±7.1 | 59.6±6.0 |
| **RF** | 15 | 64.1±2.9 | 60.8±1.3 | 46.8±3.2 | 40.5±7.1 |
|  | 20 | 71.5±7.9 | 56.2±1.3 | 43.7±4.0 | 46.5±3.9 |
|  | 30 | 78.1±6.2 | 63.9±4.2 | 55.5±3.6 | 52.0±6.0 |
|  | 45 | 81.0±5.1 | 65.4±4.3 | 60.7±6.0 | 50.6±3.7 |
| **ST** | 15 | 46.9±2.1 | 43.9±1.7 | 33.1±2.8 | 28.6±4.3 |
|  | 20 | 49.4±4.2 | 39.5±0.7 | 31.2±1.9 | 30.5±2.0 |
|  | 30 | 58.4±4.0 | 42.7±4.7 | 38.9±2.4 | 34.4±5.0 |
|  | 45 | 55.6±4.6 | 43.6±4.7 | 41.8±4.7 | 33.2±3.4 |
| **TA** | 15 | 39.8±1.8 | 36.9±1.9 | 29.4±2.8 | 26.2±2.8 |
|  | 20 | 41.0±3.1 | 32.7±2.1 | 27.0±2.7 | 28.3±1.4 |
|  | 30 | 44.9±4.0 | 37.3±0.9 | 34.7±3.7 | 33.1±2.5 |
|  | 45 | 43.5±1.5 | 36.5±2.2 | 35.0±2.9 | 31.5±3.2 |
| **QF** | 15 | 37.6±2.7 | 33.8±2.8 | 25.9±1.5 | 24.1±3.7 |
|  | 20 | 35.4±3.0 | 31.9±4.2 | 23.7±1.6 | 24.3±3.9 |
|  | 30 | 40.8±1.9 | 32.3±5.2 | 30.4±3.7 | 29.2±1.6 |
|  | 45 | 42.1±3.9 | 35.7±3.1 | 34.3±3.0 | 28.1±4.8 |
| **AM** | 15 | 28.6±3.7 | 24.0±0.9 | 20.5±2.1 | 19.3±5.6 |
|  | 20 | 24.7±2.5 | 21.1±4.3 | 18.2±1.9 | 20.0±2.6 |
|  | 30 | 33.0±3.6 | 29.0±3.1 | 26.0±1.9 | 24.0±2.8 |
|  | 45 | 33.3±2.0 | 28.6±3.4 | 28.7±3.7 | 23.1±3.5 |
| **FDL** | 15 | 28.7±3.1 | 25.3±1.0 | 20.2±0.6 | 19.2±4.8 |
|  | 20 | 28.6±1.4 | 23.6±2.5 | 18.8±1.5 | 20.0±1.0 |
|  | 30 | 31.9±2.9 | 25.9±2.2 | 23.5±1.3 | 22.5±1.8 |
|  | 45 | 33.7±2.7 | 28.6±2.2 | 25.4±2.5 | 22.6±1.6 |
| **BFA** | 15 | 21.6±3.1 | 20.0±2.4 | 16.5±1.5 | 13.7±3.2 |
|  | 20 | 22.5±4.1 | 19.2±0.2 | 14.4±1.0 | 15.7±0.5 |
|  | 30 | 26.6±3.5 | 21.5±1.8 | 20.0±2.9 | 19.0±3.0 |
|  | 45 | 28.0±4.2 | 21.7±3.4 | 22.7±2.3 | 19.0±2.6 |
| **CA** | 15 | 22.2±2.1 | 21.0±1.8 | 16.9±1.9 | 14.7±2.7 |
|  | 20 | 22.1±5.0 | 17.6±1.6 | 14.8±1.3 | 16.1±1.7 |
|  | 30 | 27.4±2.7 | 20.5±1.3 | 18.9±1.9 | 18.1±2.3 |
|  | 45 | 25.4±1.9 | 21.4±1.8 | 20.5±1.1 | 18.2±1.8 |
| **AL** | 15 | 18.8±4.1 | 16.3±1.9 | 12.8±0.9 | 11.2±3.2 |
|  | 20 | 15.1±3.4 | 15.4±4.5 | 11.9±1.1 | 11.9±0.3 |
|  | 30 | 20.5±2.4 | 14.9±1.5 | 14.1±1.3 | 13.4±1.5 |
|  | 45 | 19.3±3.3 | 17.2±2.1 | 17.1±2.4 | 13.5±2.1 |
| **PLANT** | 15 | 16.4±0.9 | 14.2±0.5 | 11.3±1.1 | 9.8±2.0 |
|  | 20 | 15.2±1.0 | 13.1±1.0 | 10.4±0.9 | 11.2±1.1 |
|  | 30 | 17.8±1.8 | 13.9±1.4 | 12.4±0.8 | 12.0±1.8 |
|  | 45 | 16.8±0.6 | 14.8±1.5 | 13.5±1.8 | 11.7±0.7 |
| **GP** | 15 | 14.5±1.7 | 11.5±0.4 | 9.6±0.8 | 8.0±1.0 |
|  | 20 | 13.5±1.2 | 9.7±1.5 | 8.1±1.2 | 8.1±0.6 |
|  | 30 | 15.1±2.4 | 11.6±1.5 | 11.5±1.2 | 10.1±1.1 |
|  | 45 | 14.0±2.2 | 12.1±2.0 | 11.8±1.3 | 9.8±1.2 |
| **GA** | 15 | 10.9±0.5 | 10.5±0.5 | 7.2±0.6 | 7.3±1.1 |
|  | 20 | 11.8±0.4 | 8.4±0.8 | 7.2±1.3 | 7.0±0.5 |
|  | 30 | 11.5±1.7 | 9.0±1.5 | 11.0±0.8 | 7.6±1.0 |
|  | 45 | 11.4±1.4 | 8.9±1.1 | 11.3±2.0 | 8.0±1.1 |
| **PL** | 15 | 10.4±0.8 | 9.8±0.9 | 8.0±0.5 | 6.7±0.9 |
|  | 20 | 10.5±1.3 | 9.2±0.5 | 7.1±0.9 | 8.2±0.5 |
|  | 30 | 11.8±1.4 | 10.3±0.9 | 9.1±0.8 | 8.5±1.1 |
|  | 45 | 12.3±1.1 | 10.4±1.2 | 9.8±1.1 | 9.0±1.0 |
| **PECT** | 15 | 10.6±0.7 | 8.4±0.2 | 6.5±1.1 | 5.6±0.9 |
|  | 20 | 10.2±0.5 | 8.5±0.2 | 6.2±0.9 | 5.7±0.9 |
|  | 30 | 11.2±1.2 | 8.8±1.3 | 8.1±1.9 | 7.3±1.6 |
|  | 45 | 12.1±1.0 | 10.2±1.4 | 9.5±1.4 | 7.6±0.8 |
| **EDL** | 15 | 8.8±0.5 | 8.2±0.6 | 6.9±0.5 | 6.1±0.8 |
|  | 20 | 9.3±1.1 | 8.2±0.4 | 5.4±0.7 | 6.1±0.4 |
|  | 30 | 9.7±0.8 | 8.7±0.8 | 7.7±0.5 | 7.5±0.9 |
|  | 45 | 9.5±1.3 | 8.4±0.9 | 8.2±0.5 | 7.4±0.8 |
| **SOL** | 15 | 7.5±0.5 | 7.1±0.7 | 6.1±0.9 | 4.7±1.1 |
|  | 20 | 7.8±0.4 | 6.2±0.4 | 4.6±0.5 | 5.7±0.6 |
|  | 30 | 7.9±1.0 | 7.3±0.5 | 6.7±1.2 | 6.3±1.1 |
|  | 45 | 7.1±0.8 | 6.7±0.4 | 7.2±1.3 | 5.9±0.7 |
| **TP** | 15 | 5.9±0.5 | 5.4±0.6 | 4.6±0.4 | 3.3±0.8 |
|  | 20 | 6.7±1.2 | 5.0±0.3 | 4.1±0.7 | 4.2±0.8 |
|  | 30 | 6.9±1.4 | 5.0±0.9 | 6.6±0.9 | 4.9±0.5 |
|  | 45 | 7.6±0.5 | 6.3±1.1 | 6.5±0.9 | 4.7±0.9 |
| **PT** | 15 | 5.6±0.7 | 4.9±0.3 | 3.3±0.2 | 2.7±1.0 |
|  | 20 | 5.5±1.5 | 4.4±0.8 | 4.0±0.6 | 3.5±0.9 |
|  | 30 | 6.4±1.0 | 5.6±1.1 | 5.2±0.5 | 4.1±0.7 |
|  | 45 | 7.1±0.2 | 5.7±0.9 | 4.7±1.0 | 4.3±0.8 |
| **POP** | 15 | 4.0±0.6 | 3.2±0.9 | 2.5±0.3 | 2.4±0.3 |
|  | 20 | 3.7±0.3 | 3.2±0.1 | 2.2±0.4 | 3.0±0.4 |
|  | 30 | 4.0±0.8 | 3.8±0.6 | 2.9±0.1 | 3.0±0.3 |
|  | 45 | 4.7±0.4 | 3.8±0.2 | 3.4±0.5 | 3.1±0.4 |

**Table S2.** Calculated % Sexual dimorphism per muscle in the mouse hindlimb over time.

|  | Age |  |  |  |
| --- | --- | --- | --- | --- |
| Muscle | P45 | P50 | P60 | P75 |
| BFA | 33.7 | 56 | 33.2 | 22.8 |
| BFP | 41.9 | 47.5 | 35.5 | 21.7 |
| ST | 43.4 | 58.5 | 50.1 | 32.7 |
| SM | 33.6 | 50.7 | 40.2 | 24.9 |
| CA | 33.2 | 49.6 | 45.1 | 24.1 |
| AM | 40.2 | 35.4 | 26.9 | 16.2 |
| AL | 51.4 | 26.6 | 45.6 | 12.7 |
| GA | 50.3 | 63.1 | 5.3 | 0.8 |
| GP | 56.4 | 65.4 | 31.6 | 18.1 |
| PECT | 61.7 | 64.5 | 37.9 | 26.4 |
| QF | 44.8 | 49.3 | 34.2 | 22.6 |
| RF | 38.7 | 63.7 | 40.7 | 33.4 |
| POP | 58.8 | 68.1 | 38.6 | 37.5 |
| GAS | 41.3 | 48.3 | 41.5 | 26 |
| PLANT | 46.4 | 45.8 | 42.6 | 24.2 |
| SOL | 25.4 | 67.9 | 18.3 | -1.2 |
| FDL | 43 | 51.7 | 35.4 | 32.8 |
| TP | 25.9 | 62.4 | 5.1 | 16.9 |
| TA | 36.6 | 51.9 | 29.4 | 24.1 |
| EDL | 29.2 | 71.1 | 26.3 | 16.5 |
| PL | 29 | 46.9 | 29.4 | 25.1 |
| PT | 64.1 | 36.8 | 24.7 | 50.1 |

Sexual size dimorphism calculated using the following formula: $[(\text{Average control male muscle weight} / \text{Average control female muscle weight}) - 1] * 100$
BFA, Biceps femoris anterior; BFP, Biceps femoris posterior; ST, Semitendinosus; SM, Semimembranosus; CA, Caudofemoralis; AM, Adductor magnus; AL, Adductor longus; GA, Gracilis anterior; GP, Gracilis posterior; PECT, Pectineus; QF, Quadratus femoris; RF, Rectus femoris; POP, Popliteus; GAS, Gastrocnemius; PLANT, Plantaris; SOL, Soleus; FDL, Flexor digitorum longus; TP, Tibialis posterior; TA, Tibialis anterior; EDL, Extensor digitorum longus; PL, Peroneus longus; PT, Peroneus tertius.

